# Phloiokeratosis – a new ichthyosiform hyperkeratotic cornification disorder in dogs with *SUV39H1* variants

**DOI:** 10.64898/2026.02.09.704839

**Authors:** Sarah Kiener, Stefan J. Rietmann, Sara Soto, Sara J. Ramos, Cherie M. Pucheu-Haston, Chi-Yen Wu, Desirae Wheatcraft, Andrew Simpson, Susanne Åhman, Brett E. Wildermuth, Michaela Drögemüller, Vidhya Jagannathan, Charles W. Bradley, Elizabeth A. Mauldin, Nadine M. Meertens, M Welle, Tosso Leeb

## Abstract

The continuous renewal of healthy epidermis depends on the finely regulated proliferation of basal keratinocytes and subsequent differentiation as the newly formed cells move upwards through the different layers of the epidermis. Perturbations in keratinocyte differentiation may lead to cornification disorders. We investigated seven dogs of different breeds belonging to four independent families that showed striking multifocal tree bark-like skin lesions. Histopathologically, lesional skin was characterized by pronounced epidermal and infundibular hyperkeratosis with epidermal and sebaceous gland hyperplasia. We therefore tentatively termed the phenotype phloiokeratosis, derived from the Greek word phloiós for tree bark and keratosis indicating abnormal keratinization. Whole genome sequencing of DNA from affected dogs revealed four independent variants in the *SUV39H1* gene encoding the SUV39H1 histone lysine methyltransferase, an H3K9 methyltransferase, which is involved in epigenetic silencing of chromatin. Phloiokeratosis is inherited as an X-chromosomal semi-dominant trait. Four of the affected dogs in our study were heterozygous females and had lesion patterns reminiscent of Blaschko lines. In two of them, trio analyses experimentally confirmed *de novo* mutation events in the *SUV39H1* gene. Previously, *Suv39h1*^*-/-*^ knockout mice had been reported to have normal skin. So far, no human patients with *SUV39H1* loss-of-function variants have been reported. The findings in *SUV39H1* mutant dogs with phloiokeratosis for the first time link SUV39H1 deficiency to a heritable skin phenotype. Our study highlights the essential role of SUV39H1-mediated epigenetic silencing during normal keratinocyte differentiation and provides a unique model for further investigations.

**Author Summary:** The integrity of the skin depends on a balanced equilibrium of keratinocyte proliferation, differentiation, and sloughing of terminally differentiated cells into the environment requiring finely regulated changes in the global transcriptome of differentiating keratinocytes. We investigated seven dogs belonging to four different families with a new disorder of cornification characterized by tree bark-like outgrowths of the epidermis. Histopathological examinations confirmed that the outermost layer of the epidermis was thickened in affected dogs. The genetic analysis yielded four different *SUV39H1* loss-of-function variants in the affected dogs from the four families. The *SUV39H1* gene encodes an enzyme that is involved in the epigenetic silencing of chromatin. The newly characterized inherited skin disease in dogs is the first clinical phenotype that has been linked to SUV39H1 deficiency. Most likely, SUV39H1 deficiency leads to delayed epigenetic silencing and consequently delayed differentiation of keratinocytes. Dogs with this rare skin disease provide an improved understanding of the essential role of SUV39H1 in the epigenetic control of gene expression in skin.

## Introduction

The skin is the largest organ in mammals. It is involved in essential functions such as temperature regulation, sensory perception, and immunologic surveillance. In addition, it provides a protective barrier against mechanical stress, dehydration, and infection [1-3]. The epidermis, which forms the outermost layer of the skin, comprises a stratified squamous epithelium that is continuously renewed by proliferating keratinocyte stem cells maintaining contact with the underlying basement membrane. The resulting keratinocytes exit the cell cycle and differentiate as they move to the upper layers of the epidermis. Ultimately, terminally differentiated cells lose their nuclei and cytoplasmic organelles to become corneocytes that form the tightly sealing stratum corneum before they are eventually sloughed into the environment. The continuous renewal of the epidermis and keratinocyte differentiation require highly coordinated changes in the expression of genes encoding cell cycle regulators, cell adhesion molecules, and structural proteins such as keratins [2,4].

On the trajectory from a stem cell to a fully differentiated keratinocyte, the number of actively transcribed genes decreases. At the same time, some specific genes, e.g., encoding specific subtypes of keratin, become very highly expressed. Epigenetic programming of chromatin plays an important role in mediating these global changes in gene expression. Basic principles of epigenetic programming have been elucidated in the last decades [5-7]. In addition to DNA cytosine methylation, a large number of histone modifications have very specific functions in activating or silencing transcription. Histone H3 lysine-9 (H3K9) methylation is an important silencing mark and characteristic of transcriptionally inactive heterochromatin [8,9]. Mammals have at least six H3K9 methylases that are able to catalyze the transfer of up to three methyl groups to H3K9 and thus initiate chromatin silencing [10,11]. These are SUV39H1, SUV39H2, EHMT1, EHMT2, SETDB1, and SETDB2. Experiments with keratinocyte cell cultures indicated that SUV39H1 and SUV39H2 in particular are involved in keratinocyte differentiation [12,13].

While the molecular mechanisms of H3K9me3 mediated chromatin silencing and the role of heterochromatin for genome stability are quite well understood, the *in vivo* physiological significance of the six different H3K9 methylases is much less clear. *Suv39h1*^*-/-*^ and *Suv39h2*^*-/-*^ knockout mice do not show any overt phenotype and have normal skin [14]. Double knockout mice lacking Suv39h1 and Suv39h2 display severely impaired viability, growth retardation and genomic instability [14]. Unlike the situation in mice, loss of function of SUV39H2 in dogs leads to hyperkeratosis of the nasal planum, while haired skin of the nose and body remains normal [15-17]. This autosomal recessive cornification disorder has been termed hereditary nasal parakeratosis (HNPK) [18,19] and results from aberrant or delayed keratinocyte differentiation [20,21]. So far, no human patients with comparable genodermatoses due to variants in genes encoding a histone methylase have been reported.

The present study was prompted by seven dogs from four different families displaying striking hyperkeratotic skin lesions. The phenotype and familial clustering suggested a genetic etiology. We therefore initiated a genetic analysis of these dogs with the aim of identifying the causal genetic variants. We report here the results of the genetic investigation while a more detailed clinical and histopathological characterization is reported separately [22].

## Results

### Clinical Phenotype

A total of 7 dogs with multifocal hyperkeratotic skin lesions were ascertained (Figure 1; S1 Table). The lesions were reminiscent of tree bark. We therefore tentatively termed the phenotype phloiokeratosis, derived from the Greek word phloiós for tree bark and keratosis indicating abnormal keratinization. The affected dogs were born between 2015 and 2024 and belong to four independent families that were numbered from A-D. Individual cases within families were designated with Arabic numbers, e.g. A-1 is the first affected dog in family A.

**Fig. 1.**
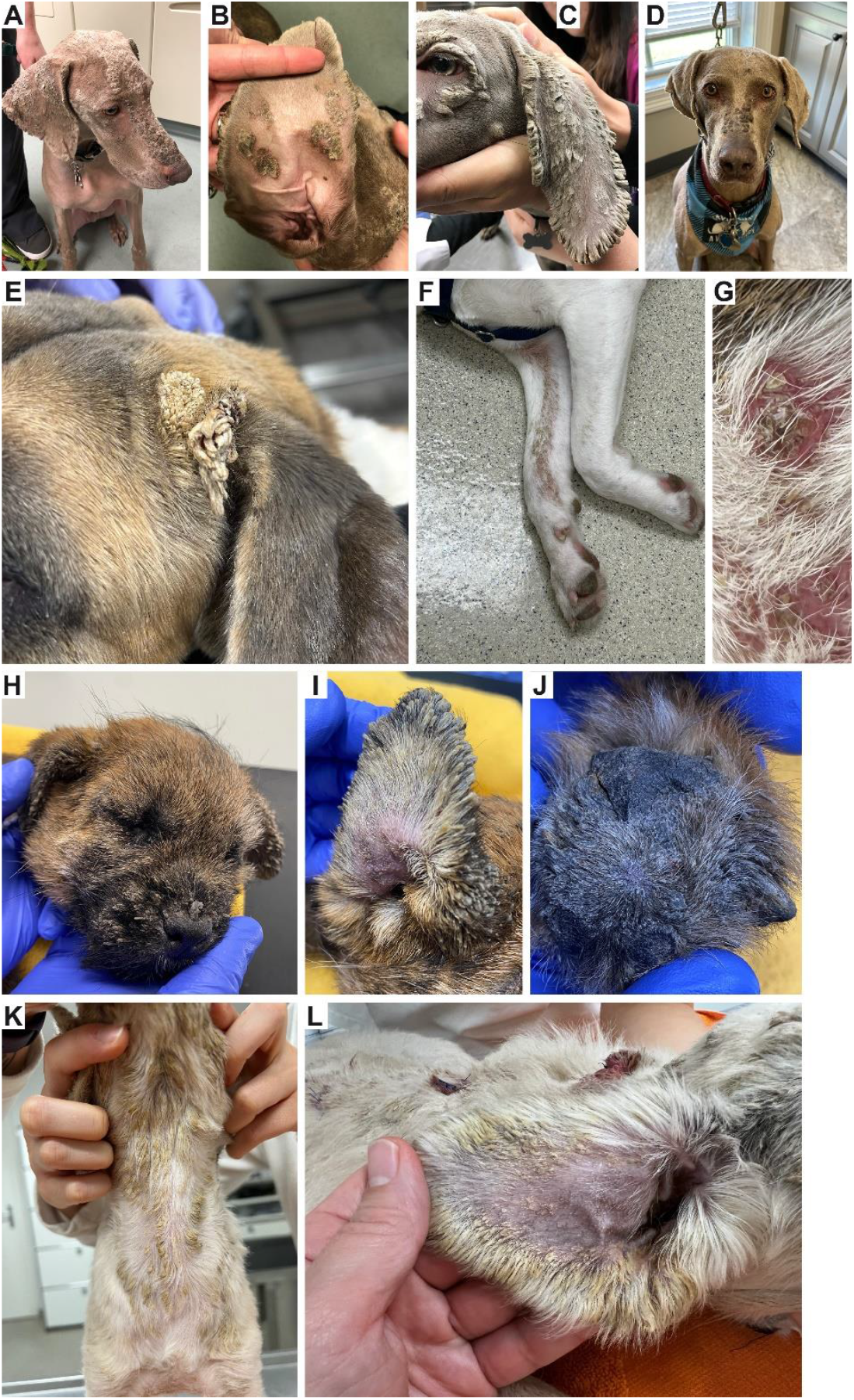
Clinical phenotype of the affected dogs. **A, B** Male Weimaraner case A-1 exhibiting severe multifocal symmetrical tree bark-like hyperkeratotic plaques on the face and ear pinnae. Less severe lesions were present at other body locations. **C** Male Weimaraner case A-2 exhibiting comparable tree bark–like hyperkeratosis on the forehead, ventral to the eyes, and convex ear pinnae, with frond-like hyperkeratosis along the ear margins. **D** Comparable lesions in case A-3, another male Weimaraner. **E, F** Female Akita mix (case B-1) showing hyperkeratotic projections at the base of the left ear and a linear hyperkeratotic lesion in a typical Blaschko pattern on the right thoracic limb. **G** Case B-2, female Akita mix, close up of a hyperkeratotic lesion on the left thoracic limb. **H-J** Female Border Terrier (case C-1) with hyperkeratotic lesions on the face, ear and paw pad. **K, L** Hyperkeratotic lesions on a Saluki (case D-1, female). Note the linear pattern of the ventral lesions in (**K**) resembling Blaschko lines.

#### Family A – Weimaraner

This family consisted of three affected male Weimaraners originating in the United States. Pedigree documents were not available. Owner reports and genetic analyses indicated that they were full siblings from two consecutive litters. Case A-1 was born in 2015, cases A-2 and A-3 were littermates born in 2016. All three dogs had moderate to severe bilaterally symmetrical, multifocal-to-coalescing hyperkeratotic plaques affecting the dorsal muzzle, periocular region, periaural region and pinnal margins. Additional lesions were noted to focally affect other areas (e.g. tip of tail, groin). Age of lesion onset was documented at 10 weeks for A-2 and 16-20 weeks for A-3. Case A-1 was rescued from a shelter. Therefore, the exact age of lesion onset is unknown in this dog. To the best of the authors’ knowledge, no other littermates were reported affected. The owner of case A-3 reported meeting the breeding parents and described no notable dermatologic abnormalities in either.

#### Family B – Akita mix

This family consisted of two affected female random-bred dogs originating in the United States that were described by their owners as Akita mixes. The affected dogs were reportedly littermates, but pedigree documents or reliable information on the parents were not available. Both dogs displayed lesions at the time of adoption at 6 months of age, primarily affecting the dorsal and lateral aspects of the trunk as well as lateral and medial aspects of the distal extremities, following Blaschko lines. B-1 developed thicker, frond-like areas of hyperkeratosis including the head and pinnae with generalized greasiness, while B-2 developed less severe lesions consisting of moderately thickened hyperkeratosis within hair follicles causing dilation and alopecia.

#### Family C – Border Terrier

Family C comprised purebred Border Terriers from Sweden. In this family, a single affected female dog together with both unaffected parents and an unaffected female littermate were examined. The affected puppy was examined at 8 weeks of age with a history of developing increasingly pronounced hyperkeratosis and scaling since a few weeks of age. At presentation there was marked hyperkeratosis affecting haired skin and paw pads. The hyperkeratosis was most pronounced in the face, around eyes and nose and along the margins of the pinnae as well as on paw pads. Generalized scaling and patchy hyperkeratosis were also present on the dorsal and lateral aspects of the body whereas the ventral aspect of the body was less affected. Both parents and a littermate were free of skin lesions.

#### Family D – Saluki

A complete litter of purebred Salukis together with their parents represented family D. The litter was born in Germany and consisted of a single affected female dog (case D-1), two healthy puppies (1 male, 1 female) and two stillborn puppies (1 male, 1 female). The parents were reportedly healthy. The affected puppy was presented with a progressive, pruritic hyperkeratosis that began at 12 days of age. Severe multifocal-to-coalescing follicular casting/hyperkeratosis was present perinasally, along the bridge of the nose, periocularly, and on the medial and lateral aspects of the pinnae. Linear hyperkeratosis following the lines of Blaschko were present on the top of the head as well as the dorsal and ventral neck. Milder lesions were present multifocally on the trunk and limbs.

### Histopathological Phenotype

Skin from lesional areas in the affected dogs was characterized by a severe mostly compact epidermal and infundibular hyperkeratosis. The hyperkeratosis was mostly orthokeratotic, but in some hyperkeratotic areas, layers of parakeratosis were present. The epidermis and the follicular epithelium were moderately to severely hyperplastic. The epidermal hyperplasia was regular. There was moderate to severe sebaceous gland hyperplasia in all affected dogs. Inflammatory cells in the dermis were scarce but prominent pigmentary incontinence was pronounced in most dogs. In biopsies from clinically non-affected skin regions, mild laminar epidermal and infundibular orthokeratotic hyperkeratosis was present histologically (Figure 2).

**Fig. 2.**
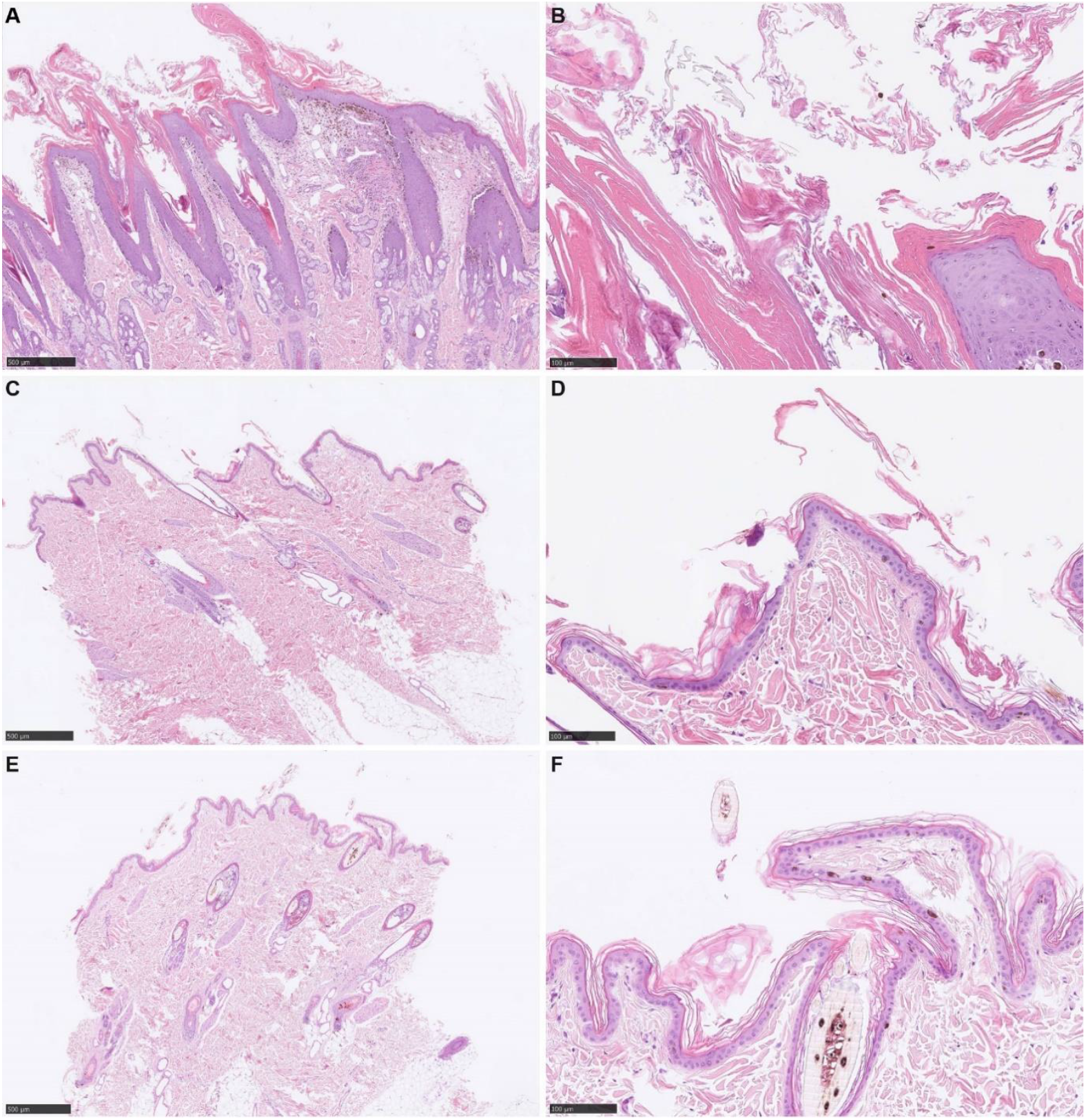
Histopathological phenotype in skin biopsies of case A-1. **A, B** Lesional skin at two different magnifications. Severe epidermal and infundibular hyperkeratosis can be seen. Sebaceous gland hyperplasia is pronounced. **C, D** Non-lesional skin of the same dog. Note the mild laminar orthokeratotic hyperkeratosis. Epidermal thickness and sebaceous glands are unremarkable. **E, F** Skin from a control Weimaraner at two different magnifications. The stratum corneum is arranged in a basket weave pattern and epidermal thickness is normal. Scale bars in A, C & E = 500 µm; B, D & F = 100 µm. Hematoxylin & eosin staining.

### Genetic Analyses

We sequenced the genomes of six affected dogs (cases A-1, A-2, A-3, B-1, C-1, D-1) and both parents of cases C-1 and D-1. In order to identify disease-associated variants, we compared the WGS data of the six cases to genomes of 1655 genetically diverse other dogs (S2 Table). Details of these analyses including all case-specific private variants are compiled in the S3 Table.

#### Family A – Weimaraner

The sequence data indicated between 66% and 70% shared haplotypes among the three affected Weimaraner dogs, corroborating the assumption that these dogs were full siblings. Given their close relationship, we hypothesized that their phenotype was due to a shared genetic variant. Filtering their whole genome sequence data for shared protein-changing variants that were absent from 1655 control genomes yielded a total of 9 candidate variants, eight heterozygous variants affecting eight different genes on autosomes and a single hemizygous variant affecting the *SUV39H1* gene on the X chromosome (Table 1, S3 Table). None of the 9 affected genes had previously been associated with a heritable skin phenotype or comparable genodermatosis as seen in the three affected Weimaraners.

**Table 1.** *SUV39H1* variants identified in dogs with hyperkeratotic lesions.

| Family | HGVS-g<br>(NC_049260.1) | HGVS-c<br>(XM_038587376.1) | HGVS-p<br>(XP_038443304.1) | Genotype in<br>affected dogs<sup>a</sup> |
| --- | --- | --- | --- | --- |
| A | g.42,103,676C>T | c.596C>T | p.(S199L) | T/- |
| B | g.42,103,621_42,103,622del | c.543_544del | p.(C181*) | TG/del |
| C | g.42,103,740_42,103,745del | c.661_666del | p.(E221_C222del) | CGAGTG/del |
| D | g.42,111,103del | c.1189del | p.(R397Vfs*75) | C/del |

<sup>a</sup>The affected dogs in family A were all male and thus had only one X chromosome. The hyphen in the genotype designation indicates the missing second allele.

As *SUV39H2* loss-of-function variants cause a related phenotype, HNPK [15,16], we considered the *SUV39H1* variant the most promising candidate. It was a missense variant, p.(S199L), predicted to affect the Pre-SET domain of the SUV39H1 H3K9-methylase (Figure 3A, B). Two different softwares yielded inconsistent predictions of the potential pathogenicity of the S199L variant. PredictSNP [23] predicted the variant as benign with a probability of 83%, while MutPred2 [24] assigned a score of 0.553, which suggests pathogenicity with low confidence.

**Fig. 3.**
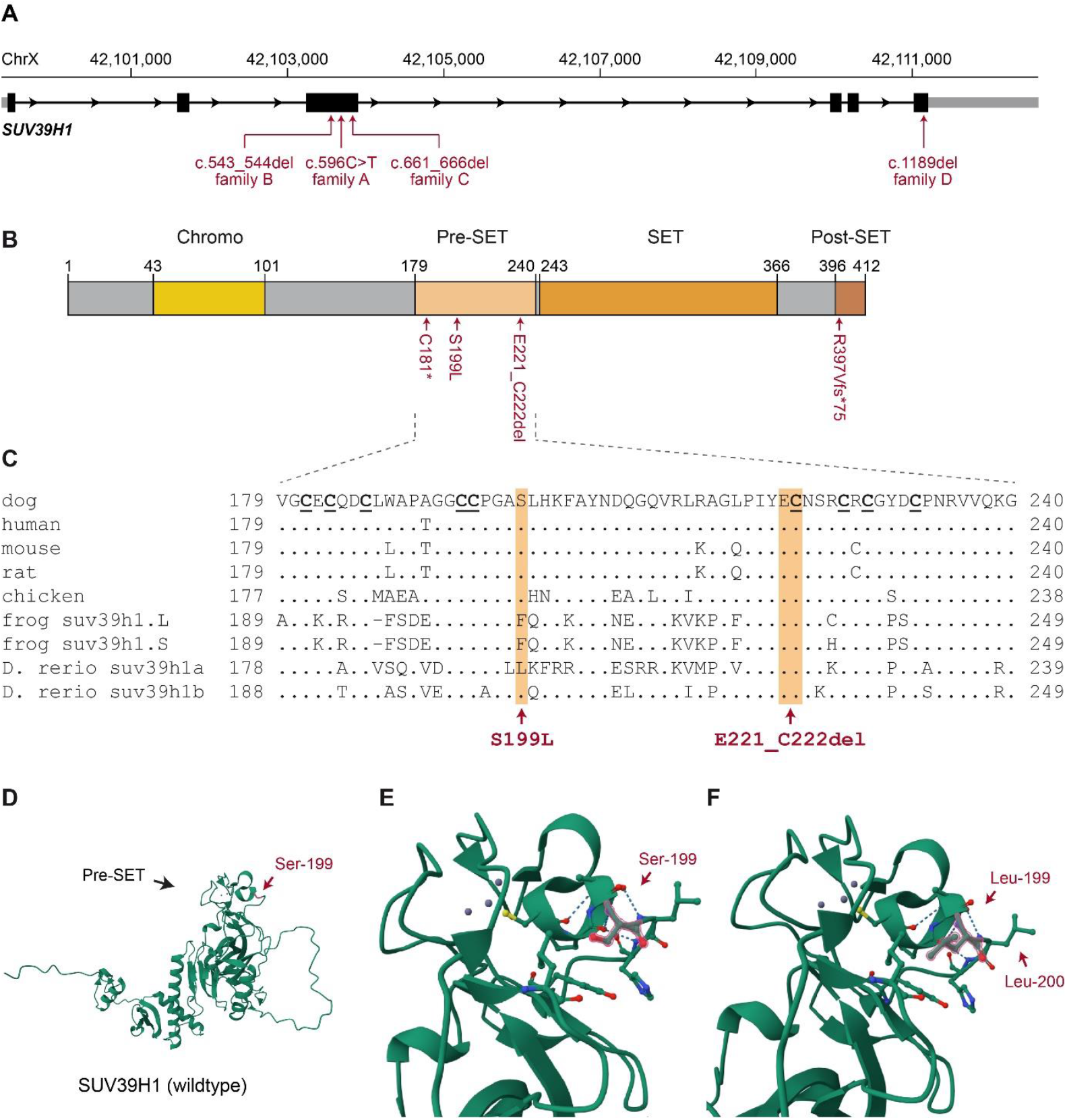
Details of SUV39H1 variants. **A** Genomic organization of the canine *SUV39H1* gene. The exons correspond to the predominant transcript isoform 2 (XM_038587376.1). Isoform 1 uses an upstream alternative first exon that is not shown. The positions of the four disease-associated variants analyzed in this study are indicated. **B** Protein domain organization of the SUV39H1 protein, isoform 2 (XP_038443304.1). Positions of the predicted effects of the four variants are indicated. **C** Evolutionary conservation of the Pre-SET domain. Nine cysteine residues that coordinate three Zn^2+^ ions are indicated in bold and underlined. The *SUV39H1* gene is duplicated in amphibians and fish. Serine-199 is conserved across mammals and birds. Glutamate-221 and cysteine-222 are strictly conserved across vertebrates. The sequences were derived from the following database accessions: *C. familiaris*, XP_038443304.1; *H. sapiens*, NP_003164.1; *M. musculus*, NP_035644.1; *R norvegicus*, NP_001389376.1; *G. gallus*, XP_040550931.1; X. laevis, NP_001084892.1 and XP_018088864.1; *D. rerio*, NP_001003592.1 and NP_001119954.1. **D** Predicted 3D-structure of wildtype canine SUV39H1. The Pre-SET domain is at the top of the molecule. **E** Enlarged detail of the Pre-SET domain with three Zn^2+^ ions indicated in grey. **F** Predicted structure of the Leu-199 mutant protein. The backbone structure is unaltered. However, with the substitution, two large hydrophobic side chains, Leu-199 and Leu-200, are in close proximity exposed at the protein surface.

SUV39H1 is evolutionarily conserved in vertebrates (Figure 3C). Due to genome duplication events, fish and amphibians have two *SUV39H1* genes whose individual functions are not known in detail. Serine-199 is strictly conserved across mammals and birds. In fish and amphibians, the residue at position 199 is not strictly conserved and e.g. the zebrafish *suv39h1a* gene has a leucine at this position, just like the Weimaraners with hyperkeratosis. This may explain why the Weimaraner p.S199L variant is predicted as benign by several of the automated classification tools.

Structural modelling indicates that serine-199 is located at the periphery of the Pre-SET domain and in contact with the aqueous solution (Figure 3D-F). It is thus interesting to note that in mammalian homologs of SUV39H1, the neighboring residue 200 is already a hydrophobic residue, while in the fish and amphibian homologs with a hydrophobic residue at position 199, the amino acid at position 200 is polar and hydrophilic. Thus, only in the S199L mutant Weimaraners, two consecutive hydrophobic residues are exposed at the protein surface.

We genotyped 131 additional Weimaraners for the p.(S199L) variant. They were all homozygous or hemizygous for the wildtype allele and none carried the mutant allele.

#### Family B – Akita mix

The genome sequence of case B-1 contained 7 homozygous and 44 heterozygous private protein-changing variants (S3 Table). As the skin lesions in this dog followed Blaschko lines (Figure 1F), we prioritized heterozygous variants on the X chromosome, of which two were present. One was a missense variant in *IL1RAPL2* encoding interleukin 1 receptor accessory protein like 2, which we did not investigate further. The other was a 2 bp coding deletion in *SUV39H1*, c.543_544del (Table 1, Figure 3A,B). The deletion shifts the reading frame and results in an immediate premature stop codon, p.(C181*). The affected sister, case B-2, was also heterozygous at this deletion.

#### Family C – Border Terrier

In this family, samples from the affected dog C-1 together with both parents and an unaffected female littermate were available for the genetic analysis. We therefore performed a trio whole genome sequencing experiment to increase the power of our variant filtering approach. The lesions in case C-1 again followed Blaschko lines and suggested a heterozygous X-chromosomal variant as underlying causal defect (Figure 1J). Thus, the most likely explanation for the clinical phenotype and family history was the occurrence of a *de novo* mutation event on the X chromosome in the maternal or paternal germline or during early embryonic development of case C-1. Analysis of the trio WGS data then indeed yielded a protein changing variant on the X chromosome that fulfilled these conditions (S3 Table). This was a 6 bp in-frame deletion in *SUV39H1*, c.661_666del (Table 1, Figure 3A,B). The deletion is predicted to remove two amino acids from the SUV39H1 protein, p.(E221_C222del). These amino acids are located in the Pre-SET domain of the enzyme, which contains 3 Zn^2+^ ions as essential cofactors. The Zn^2+^ ions are coordinated by 9 cysteine residues in the Pre-SET domain and cysteine-222 is one of them (Fig. 3C). Cysteine-222 is strictly conserved in all vertebrate SUV39H1 homologs examined. It is thus conceivable that the deletion of cysteine-222 and its neighboring amino acid will lead to a major disruption of the Pre-SET domain and thus SUV39H1 function.

Analysis of the trio whole genome sequencing data confirmed correct parentage and the *de novo* mutation event as the mutant allele was not present in the sequence data of either sire or dam. We genotyped all available members of the family by PCR and Sanger sequencing and verified that the mutant allele is exclusively present in the affected dog (Figure 4C).

**Fig. 4.**
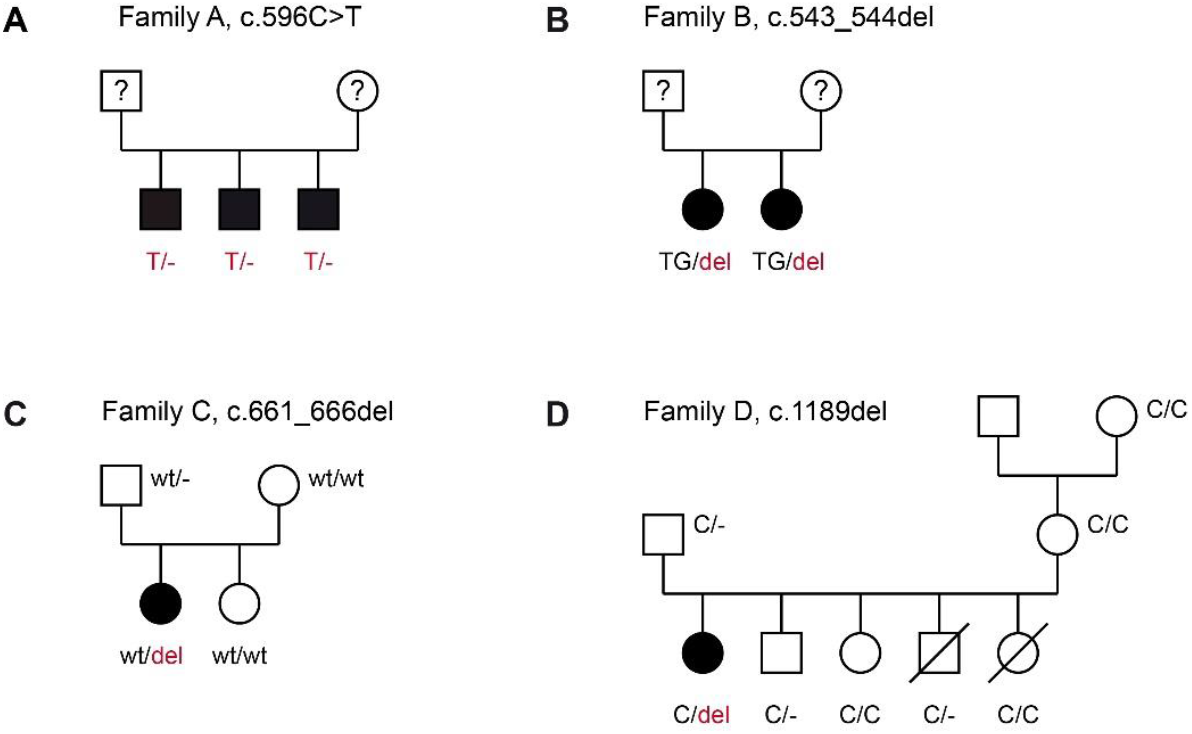
Pedigrees and *SUV39H1* genotypes of families A-D. **A** Weimaraner family with 3 affected male offspring. The phenotypes and genotypes of the parents were considered unknown as they were not examined by a veterinary dermatologist. According to one puppy owner, the skin of the parents was unremarkable. **B** Akita mix family with two affected female offspring. The phenotypes and genotypes of the parents are unknown. **C** Border Terrier family. Genotypes were determined on EDTA blood samples. Note that the deletion allele in the affected dog must have arisen due to a *de novo* mutation event, either in the maternal or paternal germline, or during early embryonic development of the affected dog. **D** Saluki family. Two stillborn puppies are indicated by strikethrough symbols. Genotypes are indicated for all dogs from which samples were available. The genotypes were determined on EDTA blood samples from the living dogs or skin tissue samples from the two stillborn puppies. Note that a *de novo* mutation event was also experimentally confirmed in family D. In the genotype designations of male dogs, the missing second allele is indicated by a hyphen.

#### Family D – Saluki

For case D-1, samples of a complete family were available for the genetic analysis. Both parents were unaffected. The lesions on case D-1 again followed Blaschko lines and suggested a heterozygous X-chromosomal variant as underlying causal defect (Figure 1K). Thus, the most likely explanation for the clinical phenotype and family history was the occurrence of another *de novo* mutation event on the X chromosome. Analysis of trio WGS data yielded a single *de novo* protein changing variant on the X chromosome, a heterozygous single base deletion in the last exon of the *SUV39H1* gene, c.1189del (Table 1, Figure 3A,B). The variant leads to a frameshift and is predicted to replace nearly the complete Post-SET domain with a much longer sequence of aberrant amino acids, p.(R397Vfs*75). Analysis of the trio whole genome sequencing data confirmed correct parentage and the *de novo* mutation event. We genotyped all available family members and verified that the mutant allele was exclusively present in the affected dog. The two stillborn littermates in this family were also clear of the mutant allele (Figure 4D).

#### Analysis of other SUV39H1 variants across 1655 control dogs

Beyond the four protein-changing *SUV39H1* variants in the seven affected dogs, only three additional protein-changing *SUV39H1* variants were present in a total of five control dogs (S4 Table). The other 1650 control dogs and the four parents of cases C-1 and D-1 did not have any protein-changing variants in *SUV39H1*.

Two of the three additional *SUV39H1* variants were missense variants predicted to affect the N-terminal region of the SUV39H1 protein, prior to the functionally important chromo domain. A male and a female Greyhound had a p.(K27M) variant in a hemizygous or heterozygous state, respectively. Both dogs had been seen as adults by a board-certified veterinary dermatologist and no signs of phloiokeratosis were present. Furthermore, a heterozygous p.(G34S) variant was present in a female Prague Ratter and a female Kunming dog. The Prague Ratter had been seen at four years of age by experienced veterinarians prior to a scheduled surgery and no evidence of phloiokeratosis was noted. The phenotype of the Kunming is unknown as the genome sequence of this dog was generated and published by another group [25].

Finally, a female Thai Ridgeback that had not shown any signs of phloiokeratosis until the age of four years was heterozygous at a nonsense variant in one of the alternative first exons. This variant is predicted to exclusively affect isoform 1 of the protein, XP_038443303.1:p.(Y6*). More details on all protein-changing variants in the canine *SUV39H1* gene are provided in the S4 Table.

## Discussion

In this study, we identified four independent *SUV39H1* variants in dogs with a new X-linked semidominant genodermatosis, tentatively termed phloiokeratosis. Phloiokeratosis clinically affects the haired skin, with or without paw pad involvement. The histopathological alterations are consistent with delayed keratinocyte differentiation and resemble the findings in dogs with *SUV39H2*-related hereditary nasal parakeratosis [15,16,18,19]. Phloiokeratosis additionally involves sebaceous gland hyperplasia. The main phenotypical difference between *SUV39H1*-related phloiokeratosis and *SUV39H2*-related hereditary nasal parakeratosis is the different anatomical localization of the lesions. For a more detailed discussion of the clinical and histopathological phenotype as well as potential therapeutic management, we refer to an accompanying manuscript [22].

The evidence for the causality of the *SUV39H1* variants differs among the families. We did not experimentally investigate the effect of the p.S199L missense variant in family A on protein function and the computational analysis of its potential effect yielded inconclusive results. However, the mutant allele was exclusively present in three affected brothers and absent from 131 breed controls as well as 1662 genetically diverse control dogs.

In family B with two affected sisters, the clinical lesions following Blaschko lines clearly pointed to functional mosaicism and an underlying heterozygous X-chromosomal variant. We initially investigated the X-chromosomal *NSDHL* gene as a potential functional candidate gene considering that heterozygous *NSDHL* variants had been reported in female dogs with linear skin lesions [26-30]. However, the WGS data of case B-1 excluded variants in *NSDHL* and revealed the heterozygous *SUV39H1*:p.C181* variant, which we assumed to cause a complete loss of SUV39H1 protein function since the coding information for more than half of the protein including the catalytic domains is lost. Unfortunately, we did not have access to the parents of the affected dogs in families A or B. We cannot exclude that the mother of the affected Weimaraners and/or one of the parents of the affected Akita mixes may already have carried the mutant allele and shown a phloiokeratosis phenotype.

The strongest evidence for causality of the *SUV39H1* variants comes from families C and D, in which we could confirm *de novo* mutation events in the affected dogs born to unaffected parents. Both variants, p.E221_C222del and p.R397Vfs*75, are predicted to disrupt functionally important parts of the protein. In our opinion, documenting two independent *de novo* mutation events in *SUV39H1*, a comparatively small gene with only six exons, leading to comparable phenotypes in dogs with different genetic backgrounds supports causality beyond any reasonable doubt. It also corroborates the findings in families A and B. A similar argumentation has been used for the identification of *MLL2* variants as the cause of human Kabuki syndrome [31]. Thus, the global collaboration of clinical specialists and the combined genetic analysis of these rare patients enabled the establishment of a completely new genotype-phenotype correlation for the *SUV39H1* gene.

The three variants in families B, C, and D were exclusively seen in heterozygous females. In contrast, the p.S199L missense variant in family A was exclusively seen in hemizygous male dogs. The currently available data are compatible with either full or partial loss of function of SUV39H1 in family A. Further research is therefore warranted to clarify whether p.S199L causes complete or only partial loss-of-function.

We identified an additional *SUV39H1* nonsense variant exclusively affecting isoform 1 in a healthy dog. This suggests that isoform 1 function might be dispensable for normal skin homeostasis, while isoform 2 is the biologically relevant isoform in this tissue.

*Suv39h1*^*-/-*^ and *Suv39h2*^*-/-*^ knockout mice do not show any overt phenotype and have normal skin [14]. In contrast, *SUV39H1*- and *SUV39H2*-mutant dogs exhibit characteristic skin phenotypes and represent the first animal models establishing cell-type specific functions of *SUV39H1* and *SUV39H2* in keratinocytes. As dogs have a special epidermis of the nasal planum that probably has no equivalent in humans, the clinical significance of *SUV39H2* function in human skin at different anatomical localizations remains unclear. However, phloiokeratosis in *SUV39H1*-mutant dogs affects normal haired skin and paw pads. It therefore remains to be seen whether humans with compromised SUV39H1 function also have a clinically relevant skin phenotype. To the best of our knowledge, no human patients with *SUV39H1* variants have been documented in the scientific literature. However, several presumable *SUV39H1* loss of function variants are reported in Gnomad [32] and several missense variants are reported as variants of uncertain significance in ClinVar [33]. Given the X-linked semidominant inheritance in dogs, a single mutant allele might cause a recognizable phenotype in humans. Conversely, a hypothetical human *SUV39H*1-related phloiokeratosis might involve a relatively mild clinical phenotype that has not been scientifically investigated so far. The results of our study in dogs suggest that *SUV39H1* should be considered a functional candidate gene in human patients with inherited cornification defects of unknown etiology. We currently do not have an explanation for the striking species differences between mice and dogs.

In conclusion, our study provides compelling evidence that *SUV39H1* loss-of-function variants cause X-linked semidominant phloiokeratosis in dogs. This establishes the first genotype-phenotype relationship for *SUV39H1*. The results imply partially overlapping roles for *SUV39H1* and *SUV39H2* in the epigenetic programming required for keratinocyte differentiation in healthy skin.

## Materials and Methods

### Ethics Statement

All examinations and animal experiments were carried out after obtaining written informed owner’s consent and in accordance with local laws, regulations, and ethical guidelines. EDTA blood samples and skin punch biopsies from affected dogs were collected for diagnostic purposes with owners’ informed consent and in accordance with ethical guidelines and approved procedures. The use of leftover material from these diagnostic specimens does not constitute an animal experiment in the legal sense. Dog blood samples from healthy control dogs were collected with the approval of the Cantonal Committee for Animal Experiments (Canton of Bern, Switzerland; permits BE 71/19 & BE94/2022).

### Animal Selection and DNA Isolation

This study included 7 dogs affected with phloiokeratosis (S1 Table), 3 unaffected relatives of case C-1 and 7 unaffected relatives of case D-1. It further included 131 control Weimaraner that were not closely related to the Weimaraner cases A-1, A-2 and A-3. Samples were processed in the VET_GEN_BERN collection of the Vetsuisse Biobank [34]. Genomic DNA was isolated from ETDA blood samples with the Maxwell RSC Whole Blood DNA Kit using a Maxwell RSC instrument (Promega, Dübendorf, Switzerland). Genomic DNA from tissue samples of the two stillborn puppies in family D was isolated with the Maxwell RSC PureFood GMO & Authentication Kit.

### Clinical and histopathological examinations

Thorough dermatological examinations on all seven affected dogs were performed by board-certified veterinary dermatologists or a board-certified veterinary dermatologist was consulted in the work-up. Each dog underwent a dermatologic evaluation that included a complete medical history, assessment of lesion distribution and morphology, and thorough examination of the skin, haircoat, claws, mucocutaneous junctions, and ears. Lesions were described using dermatologic terminology, and clinical findings were documented at each visit. Skin biopsies for histopathology were obtained from representative lesions in six of the seven affected dogs (A-1, A-2, A-3, B-1, C-1, D-1). Biopsy sites were selected to include primary lesions when possible. Sedation and local anesthesia were administered as needed, and multiple punch biopsies were collected using appropriately sized biopsy punches. Samples were gently handled to avoid crush artifact, immediately fixed in 10% neutral buffered formalin, and submitted for histopathologic evaluation by board-certified veterinary pathologists. The formalin-fixed biopsies were embedded in paraffin, cut as 4-µm sections and stained with hematoxylin and eosin (HE) prior to histological evaluation. Additional periodic acid-Schiff (PAS) stain was performed to exclude Malassezia yeasts and/or fungal and yeast infections.

### Whole Genome Sequencing and Variant Calling

For whole genome sequencing, Illumina TruSeq PCR-free DNA libraries with ∼400 bp insert size were prepared and sequenced on a NovaSeq 6000 instrument. Coverage for the 6 affected dogs ranged from 26x to 42x, and coverage for the four unaffected parents ranged from 22x to 27x. The reads were mapped to the UU_Cfam_GSD_1.0 reference genome assembly and single nucleotide variants and small indels were called using GATK HaplotypeCaller [35] as previously described [36]. The SnpEff software together with NCBI annotation release 106 was used to predict the functional effects of the called variants [37]. The genome sequences were deposited with the European Nucleotide Archive and accession numbers are given in the S2 Table.

### Parentage verification and molecular analyses of relationships

Genotypes for parentage verification were either directly obtained by genotyping DNA samples on illumina canine_HD microarrays comprising 220,853 markers or by extracting the corresponding genotypes from existing whole genome sequence data. Relationships were then investigated by performing pairwise IBD estimations with the Plink software [38].

### Sanger Sequencing

Candidate variants were confirmed and genotyped in additional dogs by generating suitable PCR amplicons followed by Sanger sequencing. PCR products were amplified from genomic DNA with AmpliTaqGold360Mastermix (Thermo Fisher Scientific, Waltham, MA, USA). Primer sequences and conditions are given in the S5 Table. After treatment of the PCR products with exonuclease I and alkaline phosphatase, the sequencing reactions were performed with ABI BigDye v3.1 kit (Thermo Fisher Scientific, Waltham, MA, USA). Sequencing products were purified by ethanol precipitation and subsequently sequenced on an ABI 3730 DNA Analyzer (Thermo Fisher Scientific, Waltham, MA, USA). Sanger sequencing data were analyzed with the Sequencher 5.1 software (GeneCodes, Ann Arbor, MI, USA).

### Protein Modeling

Protein modeling was done with AlphaFold2 via Google Colab [39,40]. Zn^2+^ ions were added to the models using AlphaFill [41]. The models were visualized using Mol* Viewer [42].

## Data availability

All data are contained in this manuscript and the supplementary files. Accession numbers for the whole genome sequence data are given in the S2 Table.

## Acknowledgements

We thank all dog owners for providing samples and information. Isabella Aebi and Carmen Rodriguez are acknowledged for expert technical assistance. We thank Philippe Plattet and Marianne Wyss for critical constructive discussions. We are grateful to the Next Generation Sequencing Platform for performing whole-genome sequencing experiments and the Interfaculty Bioinformatics Unit of the University of Bern for providing the computational infrastructure. This study was funded in part by grant 310030_200354 from the Swiss National Science Foundation.

## Author contributions

Sarah Kiener investigation, visualization, writing – original draft, writing – review & editing

Stefan J. Rietmann investigation, writing – review & editing

Sara Soto investigation, visualization, writing – original draft, writing – review & editing

Sara J. Ramos investigation, visualization, writing – review & editing

Cherie M. Pucheu-Haston supervision, writing – review & editing

Chi-Yen Wu investigation, visualization, writing – original draft, writing – review & editing

Desirae Wheatcraft investigation, visualization, writing – review & editing

Andrew Simpson investigation, visualization, writing – review & editing

Susanne Åhman investigation, visualization, writing – review & editing

Brett E. Wildermuth investigation, visualization, writing – review & editing

Michaela Drögemüller resources, writing – review & editing

Vidhya Jagannathan data curation, writing – review & editing

Charles W. Bradley investigation, writing – review & editing

Elizabeth Mauldin investigation, writing – review & editing

Nadine M. Meertens investigation, writing – review & editing

Monika Welle investigation, writing – original draft, writing – review & editing

Tosso Leeb conceptualization, supervision, funding acquisition, investigation, visualization, writing – original draft, writing – review & editing

## Supplementary information

**S1 Table**. Compilation of 7 *SUV39H1* mutant dogs with phloiokeratosis

**S2 Table**. Accession numbers of whole genome sequences from 1665 dogs

**S3 Table**. Results of variant filtering in genome sequences of 6 affected dogs

**S4 Table**. Curated protein-changing variants in the canine *SUV39H1* gene

**S5 Table**. PCR primer sequences for genotyping canine *SUV39H1* variants

